# Ubiquitination of secretory granules promotes crinophagic degradation in *Drosophila*

**DOI:** 10.1101/2025.09.24.678350

**Authors:** Tamás Csizmadia, Anna Dósa, Asha Kiran Maddali, András Jipa, Hajnalka Laczkó-Dobos, Péter Lőw, Gábor Juhász

## Abstract

Gland cells dynamically regulate their secretory granule content via balancing the rates of synthesis, maturation, secretion, and lysosomal degradation (crinophagy). The signal(s) leading to crinophagic breakdown of secretory granules are unknown. Here we show that dynamic ubiquitination of unreleased or low-grade glue-containing secretory granules marks these vesicles for crinophagy in larval salivary gland cells of *Drosophila*. We identify the ubiquitin ligase Cnot4 and the deubiquitinating enzyme Usp7 as mediators of glue granule ubiquitination and deubiquitination, respectively. Loss of either *Cnot4* or *Usp7* impairs glue granule fusion with lysosomes. Overexpression of Cnot4 induces premature crinophagy while Usp7 overexpression prevents developmental crinophagy via modulation of glue granule ubiquitination status. Our work establishes that ubiquitination of secretory granules is a key trigger of crinophagy in *Drosophila*, paving the way for further analysis of this barely characterized degradation route in Metazoans.

## INTRODUCTION

Secretory mechanisms are essential components of numerous fundamental biological processes, including digestion, respiration, neuronal and endocrine functions in all animals and humans. Secretory granules serve as storage organelles for secretory materials, and these undergo complex intracellular processes including growth through homotypic fusion, maturation via moderate acidification, and fusion with elements of the endo-lysosomal system [1–3]. The maturation process alters the composition and pH of the secretory cargo, which help prevent premature secretion and degradation of secretory granules, implying that crinophagy functions as a secretory granule quality control mechanism [3–7]. Consequently, mature secretory granules are prepared for releasing their contents through exocytosis. Notably, the effective release of this secretory material necessitates the coordinated recruitment and function of a secretion-specific acto-myosin system on the cytosolic surface of the secretory granule membrane, which already formed a fusion pore via merging with the plasma membrane, as observed in the *Drosophila* larval salivary glands and in the rat exocrine pancreas cells [8,9].

The limited secretory activity of cells can result in the accumulation of unreleased secretory granules within the cytoplasm, which are frequently degraded through crinophagy. This process, although less well-known, represents an unconventional autophagic mechanism that occurs in all secretory tissues, including exocrine, endocrine, and neuroendocrine cells [10]. Crinophagy refers to the direct fusion of superfluous secretory granules with late endosomes-lysosomes, thereby integrating the secretory cargo into the endo-lysosomal system. This process results in intense acidification, digestion, and recycling of unused secretory material [10–12]. Consequently, unreleased degradable granules transform into crinosomes, specialized secondary lysosomal compartments where the secretory contents are loosened, degraded, and recycled [11,13]. Although crinophagy was initially identified with electron microscopic examination by Smith and Farquhar in 1966 [12], the molecular mechanisms and genetic regulation underlying this process are just beginning to unfold.

Early ultrastructural investigations in the late larval salivary gland of *Drosophila pseudoobscura* identified crinophagic degradation of mucopolysaccharide (glue)-containing secretory granules [14]. Similarly, the fusion of glue granules with late endosomes or lysosomes in the late larval and prepupal salivary glands of *Drosophila melanogaster* is also developmentally programmed. As a result, this organ provides a robust experimental system for investigating the intricate molecular mechanisms and regulatory pathways that control crinophagy [7,11]. In our previous investigation, we developed fluorescent reporter systems to monitor the fusion of secretory granules and late endosomes-lysosomes, and identified the key components involved in secretory granule-lysosome fusion within late larval salivary gland cells of *Drosophila* [11,15,16]. However, the molecular signal(s) that trigger the selective crinophagic degradation of obsolete or low-quality secretory granules instead of exocytosis are still unknown.

The human body also contains several protein secreting gland tissues, all of which utilize the process of crinophagy to regulate their secretory granule pool. This mechanism eliminates unnecessary secretory vesicles and ensures quality control of the secretory material [7,10,12,17,18]. In the exocrine pancreas, the extensive fusion of trypsinogen granules with lysosomes may trigger the premature intracellular activation of trypsinogen by lysosomal hydrolases such as cathepsins, leading to necrotic cell death of the gland cells and severe inflammation in the pancreas [19,20]. Crinophagy is also believed to modulate the amount and production of insulin in the β-cells of the Langerhans islets [17,21–23]. In conclusion, a deeper understanding of the genetic network governing the process of crinophagy is crucial for improving our knowledge and potential prevention of acute pancreatitis and abnormal insulin production observed in type 2 diabetes.

Ubiquitin is a highly conserved protein and serves as a posttranslational modifier, playing diverse roles in cellular processes. This small molecule can be covalently conjugated to diverse target biomolecules, including proteins, lipids, and carbohydrates, in the form of mono-, multi, or polyubiquitin chains that employ various linkage methods, such as Lysine-48 (K48) and Lysine-63 (K63) types [24–26]. Eukaryotic cells exhibit several examples of ubiquitination and selective macroautophagic degradation of damaged or obsolete cellular organelles, such as macromitophagy, lysophagy, and macrosecretophagy, all of which necessitate autophagosome formation [7,20,27]. Components of the ubiquitin-proteasome system, including E3 enzymes or deubiquitinating enzymes (DUBs), are also critical for vesicular trafficking and autophagic pathways [24,28,29]. Intriguingly, in yeast cells, the vacuolar import and degradation (Vid) pathway functions as a specialized, autophagosome-independent autophagic process akin to crinophagy, and this mechanism involves the ubiquitin ligase encoding gene *Vid24/YBR105C* [11,30].

In this study, we examined the exciting question of how secretory granules that are unnecessary or of low-quality are directed to the lysosomal compartment. We discovered that ubiquitin serves as a molecular signal that localizes to the surface of glue-containing secretory granules when crinophagy is developmentally activated, guiding them for degradation in the late larval salivary gland of *Drosophila*. Additionally, we identified two novel regulators of this process: the ubiquitin ligase Cnot4, which we find to be required for ubiquitin coating of glue granules, and the deubiquitinase Usp7, the primary factor responsible for removing ubiquitin from these granules during normal development. Strikingly, the silencing of either of these genes resulted in defects in glue granule acidification and granule-to-lysosome fusion, similar to previously identified essential crinophagic factors, such as *Vps16A*, *Rab7*, and *Syx13* [11]. Our findings provide new insights into the regulatory mechanism of crinophagy and open a new avenue for functional analysis of this process in animals and humans.

## RESULTS AND DISCUSSION

### 1. Ubiquitin is recruited to the surface of glue granules at the onset of developmentally programmed crinophagy

Based on previous studies on the role of ubiquitin in the macroautophagic degradation of various cellular organelles, such as mitochondria, chloroplasts, peroxisomes, and small secretory vesicles [20,24,27], and vid vesicle-vacuole fusion in yeast [30], we hypothesized that the crinophagic breakdown of the large, glue-containing secretory granules in the late larval salivary gland cells of *Drosophila* may also be ubiquitin-dependent. To test the localization of ubiquitin on secretory granules, we analyzed salivary glands from a transgenic *Drosophila* stock that simultaneously expressed glue-DsRed to label glue-containing secretory granules and GFP-tagged ubiquitin (GFP-Ub), which we refer to us the glue-ubiquitin reporter assay [31,32]. This allowed us to examine the localization of these reporter proteins during the naturally activated crinophagy in the late larval and prepupal salivary gland cells. During the wandering developmental phase (-6 h RPF – relative to puparium formation), the salivary gland cells exhibited numerous glue-DsRed positive granules, and GFP-ubiquitin was uniformly detected throughout the cytoplasm and the nucleus, excluding the nucleolus (**Figure 1A and I**). During the late larval stage, when developmentally programmed crinophagy is typically activated in salivary gland cells [11], there was a pronounced and evident recruitment of GFP-ubiquitin onto the surface of numerous granules containing glue-DsRed (yellow arrowheads, **Figure 1B and I**). The salivary glands of prepupal animals (0 h RPF) already exhibited weaker GFP-ubiquitin signal on the membrane of certain glue granules-crinosomes (pale yellow arrowheads on **Figure 1C and I**). These findings reveal a robust association between the developmental stage characterized by elevated ubiquitin levels on the glue granules and the commencement of the naturally triggered, intense fusion process between the glue granules and lysosomes (happening at -2 h RPF) [11]. The intensity of GFP-ubiquitin on glue granules-crinosomes decreased in the salivary gland cells of prepupal animals, suggesting the removal of ubiquitin from the membrane of glue containing structures. These findings raise the possibility that ubiquitin functions as a marker to target secretory granules toward degradation, possibly acting as a signal to initiate crinophagy. Furthermore, our results indicate that ubiquitin only transiently associates with glue granule-crinosomal membranes, implying a sophisticated regulatory mechanism connected to the potential fates of glue granules in the late larval salivary gland cells.

**Figure 1.**
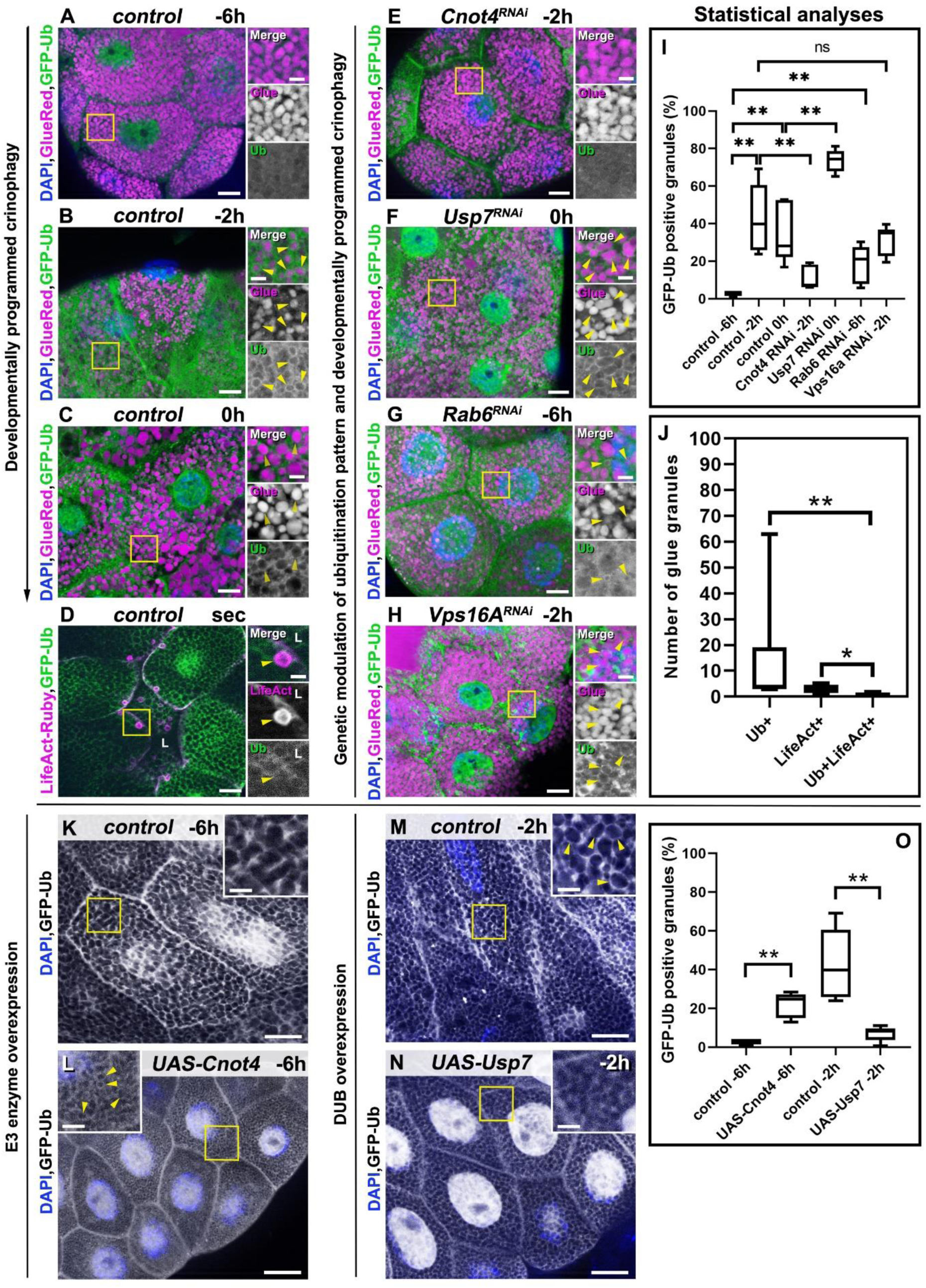
Dynamic ubiquitination of glue granules during crinophagy depends on Cnot4 and it is antagonized by Usp7. (**A-C**) Time course of ubiquitin localization during the naturally activated (developmentally programmed) crinophagy in the *Drosophila* larval and prepupal salivary gland cells coexpressing glue-DsRed and GFP tagged ubiquitin (GFP-Ub). (**A**) Salivary gland cells from wandering larvae (-6h RPF) contain many DsRed positive glue granules, and GFP-Ub is seen in the cytoplasm and in the nucleus but not in the nucleolus. (**B**) In the late larval stage, 2h before puparium formation (-2h RPF), GFP-Ub localizes on the membrane of multiple glue granules (yellow arrowheads) in the salivary gland cells (**C**). A faint signal of GFP-Ub is still visible on the membrane of glue granules-crinosomes (yellow arrowheads) in the salivary glands from white prepupae (0h RPF). (**D**) Ubiquitin does not colocalize with the exocytotic marker LifeAct-Ruby on glue granules in the salivary gland cells from the secretion period, co-expressing Life-Act-Ruby and GFP-Ub. The large LifeAct-Ruby positive structures are glue granules undergoing exocytosis [8,9], but they exhibit not too low GFP-Ub signal (yellow arrowheads). Sec: secretion period, L: lumen. (**E-H**) Silencing of selected genes in the *Drosophila* larval and prepupal salivary gland cells expressing glue-DsRed and GFP-Ub. (**E**) Knockdown of *Cnot4* prevents ubiquitination of glue granules in late larvae (-2h RPF), they appear similar to gland cells from younger animals (-6h RPF - **A**). (**F**) Silencing of *Usp7* results in persisting, strong GFP-Ub signal on the membrane of glue granules-crinosomes (yellow arrowheads) in prepupae (0h APF) compared to the control (**C**) cells. (**G**) Silencing of *Rab6* triggers premature recruitment of GFP-Ub onto the membrane of immature glue granules (yellow arrowheads) in the salivary gland cells from the wandering animals (-6h RPF), unlike the control (**A**). (**H**) Knockdown of *Vps16A* does not disrupt the ubiquitin-positivity of the membrane of glue granules (yellow arrowheads) in late larval salivary gland cells (-2h RPF). (**I**) Quantification of data from panels A-C and E-H, n=6-8 animals, Mann-Whitney tests. P values are ** p<0.0022 and ns=0.3939. (**J**) Quantification of data of panel D, n=6-8 animals, Mann-Whitney tests. P values are **p<0.0022 and *p<0.0108. The boxed regions in panels (**A–H**) are shown as enlarged insets on the right side of each panel. (**K-N**) Overexpression of Cnot4 or Usp7 perturbs GFP-Ub localization on the surface of glue granules. Salivary gland cells from wandering animals (-6h RPF) co-expressing GFP-Ub and Cnot4 already contain GFP-ubiquitin positive glue granules (**L**, yellow arrowheads), compared to the control cells (**K**). A granule ubiquitination defect is seen in Usp7 overexpressing salivary gland cells (**N**) from late larvae (-2h RPF), compared to the control cells (**M,** yellow arrow-heads). Quantification of data from panels K-N (**O**) was performed on samples from 6-8 animals using Mann-Whitney tests. P values are ** p<0.0022 (**O**). Magenta and green channels of merged images are also shown separately as indicated. The boxed regions in panels (**K–N**) are shown enlarged. Scale bars of A–H and K-N panels 20 µm, insets 5 µm.

### 2. Ubiquitin does not colocalize with the exocytotic marker LifeAct-Ruby on the membrane of glue granules

The molecular composition of ubiquitin modifications on organelles can induce diverse cellular processes, such as the initiation of autophagic degradation or the regulation of vesicular transport pathways [26]. The possible fates of glue-containing secretory granules after their maturation may be exocytosis or crinophagic degradation through intense late endosomal-lysosomal fusions. We wanted to investigate the role of ubiquitin on the membrane of glue granules, so we next tested whether ubiquitin takes part in the designation of glue granules to the exocytotic pathway. To this end, we examined secreting salivary glands that simultaneously expressed GFP-ubiquitin and LifeAct-Ruby, the marker of secretory granule-apical membrane fusion [8,9] and we observed numerous LifeAct-Ruby-positive granular structures localized near the apical membranes of the cells, which were associated with the expanding glandular lumen (**Figure 1D**). Of note, GFP-ubiquitin did not colocalize with LifeAct-Ruby on the surface of the glue granules undergoing secretion, suggesting that ubiquitin does not play a prominent role in the exocytosis of these granules (**Figure 1D and J**).

### 3. The RING-type ubiquitin ligase Cnot4 and the deubiquitinating enzyme Usp7 are required for the normal dynamics of glue granule ubiquitination during the late larval-prepupal transition of *Drosophila*

Ubiquitination, a dynamic post-translational modification, is carried out by specialized enzymes. The activity of E3 enzymes facilitates the attachment and assembly of diverse ubiquitin patterns on the substrate molecule, while deubiquitinases perform the opposite function [24]. Motivated by the robust GFP-ubiquitin labeling of glue granules in salivary gland cells during the late larval developmental stage, we conducted a targeted knockdown screen in transgenic *Drosophila* flies that co-expressed glue-GFP and glue-DsRed to investigate crinophagy based on the quenching of GFP (but not DsRed) within the acidic lumen of lysosomes [11]. Positive controls in our screen included *dor*, *lt*, and *Vps11*, genes encoding RING domain-containing subunits of the HOPS vesicle tethering complex, whose knockdown inhibited crinophagy as expected [11]. From the several potential new hits (**Table S1-2**), we decided to study the crinophagic role of the genes *Cnot4* (which encodes a RING - Really Interesting New Gene - type E3 enzyme), and the deubiquitinase *Usp7*. We chose to focus on these two genes because their knockdown phenotypes were the most robust without perturbing the size and number of glue granules, and no gland atrophy was observed either. We thus further investigated the roles of *Cnot4* and *Usp7* using our newly developed glue-ubiquitin reporter system.

Cnot4 (CCR4-NOT transcription complex, subunit 4) is an E3 enzyme that is a subunit of the CCR4-NOT (Carbon Catabolite Repression 4 - Negative On TATA-less) deadenylase complex involved in mRNA degradation [33]. Interestingly, in *Drosophila* Cnot4 is not stably incorporated into the CCR4-NOT complex, suggesting that it may possess additional independent functions [34]. Silencing the *Drosophila Cnot4* gene in late larval salivary gland cells resulted in glue granules lacking the GFP-ubiquitin signal (**Figure 1E and I**), unlike control cells at a similar developmental stage (**Figure 1B**). Furthermore, overexpression of the Cnot4 protein in the salivary gland cells of wandering larvae led to premature ubiquitination of numerous glue granules (**Figure 1L and O**), in comparison to control cells (**Figure 1A, K and O**). This observation indicates that Cnot4 plays a crucial role in the ubiquitination of glue granule membranes during developmentally programmed crinophagy in salivary gland cells.

Usp7 is a deubiquitinating enzyme that performs several key cellular functions, such as regulating tumorigenesis [35], modulating endosomal recycling pathways [36], and protecting the lysosomal Transcription Factor EB (TFEB) from proteasomal degradation [37]. Silencing this gene specifically in salivary gland cells resulted in an effect opposite to *Cnot4* RNAi (**Figure 1E**): we observed persistent, strong GFP-ubiquitin signals on the surface of glue granules-crinosomes in prepupal salivary gland cells (0h APF, **Figure 1F and I**), in contrast to the faint GFP-ubiquitin signals seen in prepupal controls (**Figure 1C**). Importantly, the overexpression of Usp7 in the salivary gland cells prevented the ubiquitination of glue granules (**Figure 1N and O**) compared to control cells (**Figure 1B, M and O**). These data suggest that Usp7 mediates the removal of ubiquitin from the membrane of glue-DsRed positive structures as glue granules transform into crinosomes.

Given the indispensable roles of Cnot4 and Usp7 in the normal ubiquitination dynamics of glue granule membrane proteins and/or lipids during the late larval-prepupal transition, ubiquitin is likely a part of a precise regulatory system that may direct glue granules towards crinophagic degradation.

### 4. Disruption of endosome-to-TGN retrograde transport causes the early ubiquitination of immature glue-containing small secretory vesicles during developmental program-independent crinophagy

In our earlier work, we employed genetic manipulation to trigger premature activation of crinophagy independent of the typical developmental process. Our findings indicated that disrupting the retrograde transport from endosomes to the trans-Golgi network (TGN) leads to premature acidification and increased crinophagic degradation of the accumulated immature and diminutive glue-containing secretory vesicles within the salivary gland cells [3]. *Rab6* is one of the genes encoding a small GTPase that plays a critical role in the endosome-to-TGN retrograde transport pathway [38]. Using our transgenic flies expressing glue-DsRed and GFP-ubiquitin, we silenced *Rab6* in larval salivary gland cells from wandering animals (-6 h RPF), which resulted in GFP-ubiquitin recruitment onto the surface of small, immature glue-DsRed positive vesicles (**Figure 1G and I**). Importantly, at this developmental stage, ubiquitin did not normally localize to the membrane of glue granules (**Figure 1A**), which implicated a potential role for ubiquitin in initiating developmental program-independent crinophagy, similar to the developmentally regulated secretory granule-lysosome fusion.

### 5. Ubiquitin is recruited to the membrane of glue granules independently of the endolysosomal system

During normal cellular processes, ubiquitin plays diverse roles in various mechanisms, including endocytosis, different types of autophagy, and the maintenance of lysosomal membrane protein homeostasis [26,28]. Ubiquitin primarily contributes to the functioning of the endo-lysosomal system, thereby maintaining cellular homeostasis. Vps16A is an essential component of this system, as it is a subunit of the CORVET (class C cORe Vacuole/Endosome Tethering - miniCORVET in *Drosophila*) and HOPS (HOmotypic fusion and Protein Sorting) tethering complexes. The CORVET complex mediates the homotypic fusion of early endosomes, while HOPS is required for homo-and heterotypical lysosomal fusion events [39]. We showed previously that Vps16A is crucial for the fusion of secretory granules to the lysosomes [11]. Since ubiquitin was shown to be recruited to the surface of endosomes and lysosomes [26,29], we sought to establish the timing of ubiquitin recruitment to the surface of glue granules during the induction of crinophagy. Therefore, we silenced *Vps16A* in the salivary gland cells of our glue-DsRed and GFP-ubiquitin expressing animals. Interestingly, we found that ubiquitin was still present on the membrane of glue granules when lysosomal fusion is inhibited by *Vps16A* silencing in the gland cells from the late larval period (-2 h RPF, **Figure 1H and I**). This observation suggests that, during the induction of crinophagy, ubiquitin is recruited directly to the surface of glue granules from the cytoplasm without the contribution of the endo-lysosomal system.

### 6. Endogenous ubiquitin is also present on the membranes of glue granules and forms K63-polyubiquitin chains

In our previous experiments, we expressed GFP-conjugated ubiquitin specifically in *Drosophila* salivary gland cells. We next used immunostaining to determine the localization of endogenous ubiquitin in the salivary gland cells expressing solely glue-DsRed during the late larval stage. Consistent with our earlier findings, endogenous ubiquitin was recruited to the membrane of glue granules in the gland cells 2 hours prior to puparium formation (-2h RPF, **Figure 2A**).

**Figure 2.**
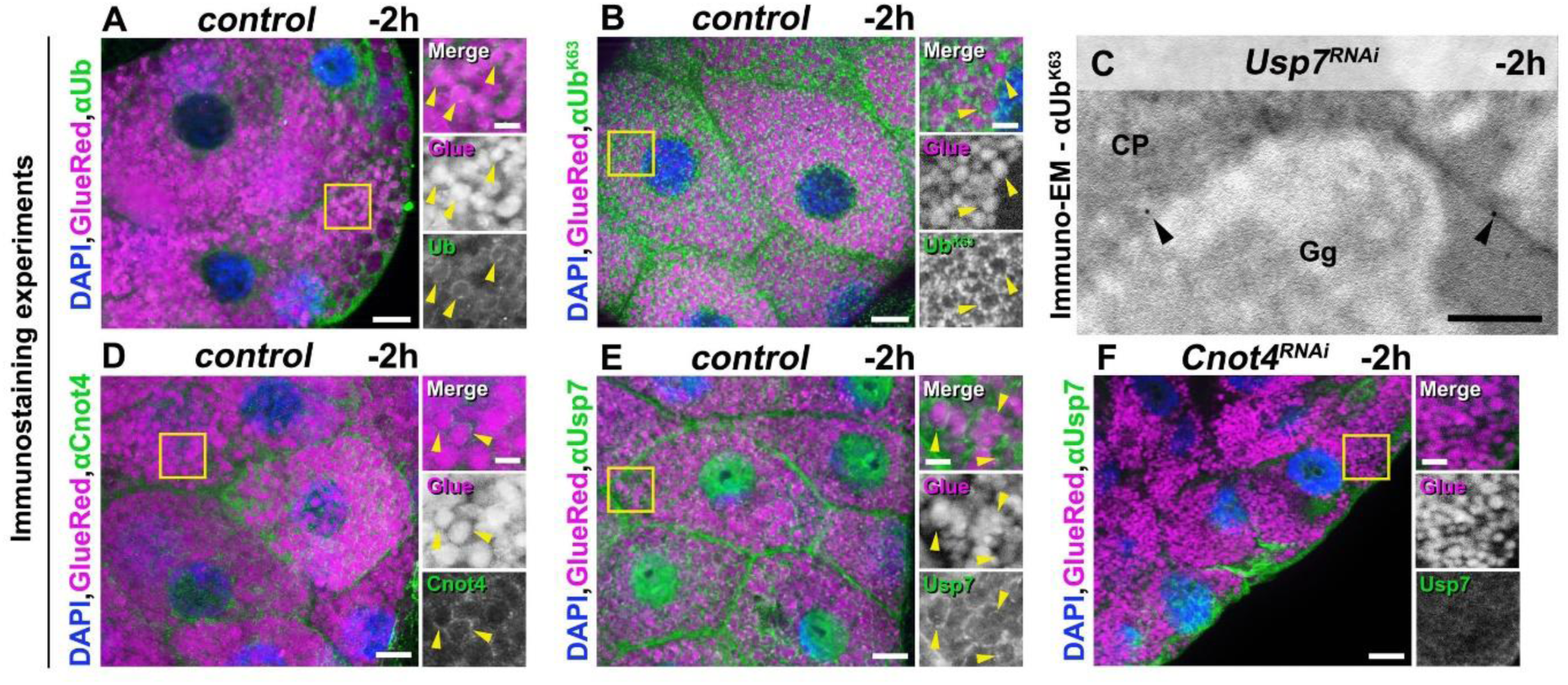
Endogenous ubiquitin, Cnot4 and Usp7 localize to the surface of glue granules. (**A-F**) Immunostaining of glue-DsRed expressing salivary gland cells from the late larval developmental stage (-2h RPF). (**A**) Immunohistochemical analysis detects endogenous ubiquitin on the surface of secretory granules (yellow arrowheads). (**B**) An antibody specific for K63linked polyubiquitin (αUb^K63^) also labels the membrane of glue granules (yellow arrowheads). (**C)** Immunogold detection by transmission electron microscope (TEM) confirms K63 linked polyubiquitin (Ub^K63^) on the surface of glue granules (black arrowheads) in *Usp7* RNAi animals. Abbreviations: CP: Cytoplasm, Gg: Glue granule. The ubiquitin ligase Cnot4 (**D**), and the deubiquitinating enzyme Usp7 (**E**) are present on the membrane of glue granules (yellow arrowheads) in the salivary gland cells from the late larval period (-2 h RPF). (**F**) Usp7 does not associate with glue granules in *Cnot4* RNAi cells (-2 h RPF). The boxed regions in panels (**A–F**) are shown as enlarged insets on the right side of each panel. Magenta and green channels of merged images are also shown separately as indicated. Scale bars of A, B and D-F panels equal 20 µm, C panel 250 nm, insets 5 µm.

The appearance of diverse ubiquitin forms on molecules and cell organelles raises the possibility that mono-, multi-, and/or polyubiquitin chains (the latter exists with various connections, such as K48, K63, K11 polyubiquitin chains) may be present on glue granules [26]. Interestingly, the maturation of secretory granules relies heavily on the endosomal system, and such vesicular trafficking and autophagic pathways are often regulated by Lysine-63 (K63)-linked polyubiquitin patterns on the surfaces of several cell organelles, such as mitochondria, endosomes, and lysosomes [26]. Based on these observations, we hypothesized that the possible type of ubiquitin found on glue granules may be K63-linked polyubiquitin. Indeed, our immunostaining experiment with a chain-specific antibody revealed that K63-linked polyubiquitin was present on glue granule membrane (**Figure 2B**). In line with this, we detected K63-linked polyubiquitin in immunogold labeling electron microscopy experiments, appearing as gold particles on glue granule membranes (**Figure 2C**). To further confirm the presence of polyubiquitin on glue granules, we immunoprecipitated GFP-ubiquitin positive structures from salivary glands containing glue-DsRed marked secretory granules. Western blot analysis using GFP (**Figure S1A**), mono- and polyubiquitin (**Figure S1B**), and DsRed (**Figure S1C**) antibodies clearly demonstrated the presence of polyubiquitinated proteins associated with glue granules in the immunoprecipitated fraction.

In conclusion, glue granules acquire K63-linked polyubiquitin chains, which are likely to serve as signals governing the vesicles direct fusion with late endosomes and lysosomes.

### 7. Cnot4 and Usp7 are recruited to the surface of glue granules during the late larval-prepupal transition

Cnot4 is a component of the CCR4-NOT complex, which plays a crucial role in mRNA turnover. Although *Drosophila* Cnot4 (dCnot4) is not stably integrated into the CCR4-NOT complex, this protein may modulate the ubiquitination of glue granule membranes either directly or through mRNA metabolism. To test this, we investigated the intracellular localization of endogenous Cnot4 using a Cnot4 antibody in *Drosophila* salivary glands. Importantly, Cnot4 was clearly observed on the surface of glue granules when developmentally programmed crinophagy was activated (-2h RPF, **Figure 2D**), suggesting that Cnot4 is a ubiquitin ligase that directly ubiquitinates the membrane of glue granules prior to their crinophagic degradation.

The deubiquitinating enzyme Usp7 also regulated glue granule ubiquitination in our experiments, so we next tested its localization. Previous research indicates that Usp7 regulates transcription by modulating Histone H2B monoubiquitination in glial cells [40], and it is also essential for the endosomal recycling pathway through regulating the actin-nucleating Wash protein [36]. In line with these, Usp7 was present at high levels in the nucleus and close to the plasma membrane of salivary gland cells (**Figure 2E**). Additionally, Usp7 can modulate the intracellular levels of the lysosomal transcription factor TFEB, influencing the auto- and heterophagic degradation pathways and lysosomal function [37]. Our immunolabeling experiment with anti-Usp7 indeed revealed the presence of this protein on the membrane of glue granules in salivary gland cells during the late larval period (**Figure 2E**). Based on this finding, we propose that Usp7 also has a direct role in the deubiquitination of glue granule membrane proteins and/or lipids.

Usp7 plays a crucial role in regulating the localization and activity of the Wash protein on the endosomal surface as part of the Trim27-containing ubiquitin ligase complex [36]. The involvement of the endosomal compartment in the maturation of glue granules [1,6,16] raises the possibility that Usp7 functions on the surface of glue granules in the late larval salivary gland cells in association with crinophagy-specific ubiquitin ligase complex(es) together with Cnot4. We thus examined whether the localization of Usp7 to glue granules would be impaired by *Cnot4* silencing. Strikingly, *Cnot4* knockdown prevented the association of Usp7 to glue granules (**Figure 2F and Figure S2E**). Usp7 recruitment thus depends on Cnot4, either directly (in complex with Cnot4) or indirectly (being recruited by Cnot4-dependent ubiquitination).

Collectively, Cnot4 and Usp7 are localized to the glue granule membrane when intense crinophagic degradation begins in these cells. Our loss of function and localization data also support that Cnot4 and Usp7 play a direct role in regulating the formation and removal of the ubiquitin pattern on the glue granule membrane, which could reroute glue granules towards crinophagic degradation.

### 8. *Cnot4* and *Usp7* are required for secretory granule-lysosome fusion and crinophagic degradation

During the late larval-prepupal developmental stage, the salivary glands undergo multiple significant morphological transformations associated with the completion of glue secretion. These changes encompass the developmentally regulated reconstruction of gland cell architecture during exocytosis, as well as the rapid removal and degradation of the remaining glue granules through extensive fusion with late endosomes and lysosomes. The initial molecular regulators identified in this process include the small GTPases Rab2, Rab7, and Arl8, the HOPS tethering complex, and the Syx13-Snap29-Vamp7/Ykt6-containing SNARE complexes [11,15,16]. Furthermore, loss of the endosome-to-TGN recycling pathway components (such as *Rab6*, *Syx16* and *Vps53*) leads to premature crinophagy in the salivary gland cells of *Drosophila* independently of the developmental program [3].

Our targeted screen (**Tables S1, S2**) indicated that *Cnot4* and *Usp7* are important for the intense acidification of glue granules and/or their fusion with lysosomes. In the acidification assay, prepupal (0 h RPF) animals co-express glue-GFP and glue-DsRed fusion proteins, which label secretory cargo-containing granules [11,32,41]. Crinosomes form in the control cells due to the fusion of glue-GFP and glue-DsRed containing secretory granules with acidic lysosomal compartments, and GFP fluorescence is rapidly quenched in the acidic crinosomal environment. Importantly, the glue-DsRed component is less sensitive to low pH, allowing the degrading cargo to retain the DsRed signal. These markers enabled us to distinguish the intact secretory granules (GFP+, DsRed+) from acidic crinosomes (DsRed+ only) [3,11,15]. Strikingly, both *Cnot4* and *Usp7* silenced salivary gland cells exhibited a strong inhibition of GFP quenching (**Figure 3B-C and J**), in contrast to the control cells that predominantly contained DsRed-positive crinosomes (**Figure 3A and J**) at the prepupal developmental stage. Importantly, the knockdown of the other components of the CCR4-CNOT complex, such as *Not1* and *Rcd-1* did not perturb the intense acidification of glue granules (**Figure S2A-C and D**). The overexpression of Cnot4 and Usp7 demonstrated contrasting outcomes regarding the acidification of glue granules. Overexpression of Cnot4 in salivary gland cells of wandering animals led to premature acidification of glue granules (**Figure S3B and I**) compared to the control cells (**Figure S3A and I**). Conversely, Usp7 overexpression induced a significant acidification defect in prepupal salivary gland cells (**Figure S3D and I**) in comparison to the control (**Figure S3C and I**). It is important to note that the acidic (only glue-DsRed positive) structures seen in **Figure 3A**, **S3B and S3C** were consistent with being crinosomes based on our ultrastructural investigations (**Figure 3G**, **S3F and G**) [11]. These crinophagy modulating effects of Cnot4 and Usp7 overexpressions are likely caused by their promoting and inhibiting of glue granule ubiquitination, respectively (**Figure 1K-N and O**).

**Figure 3.**
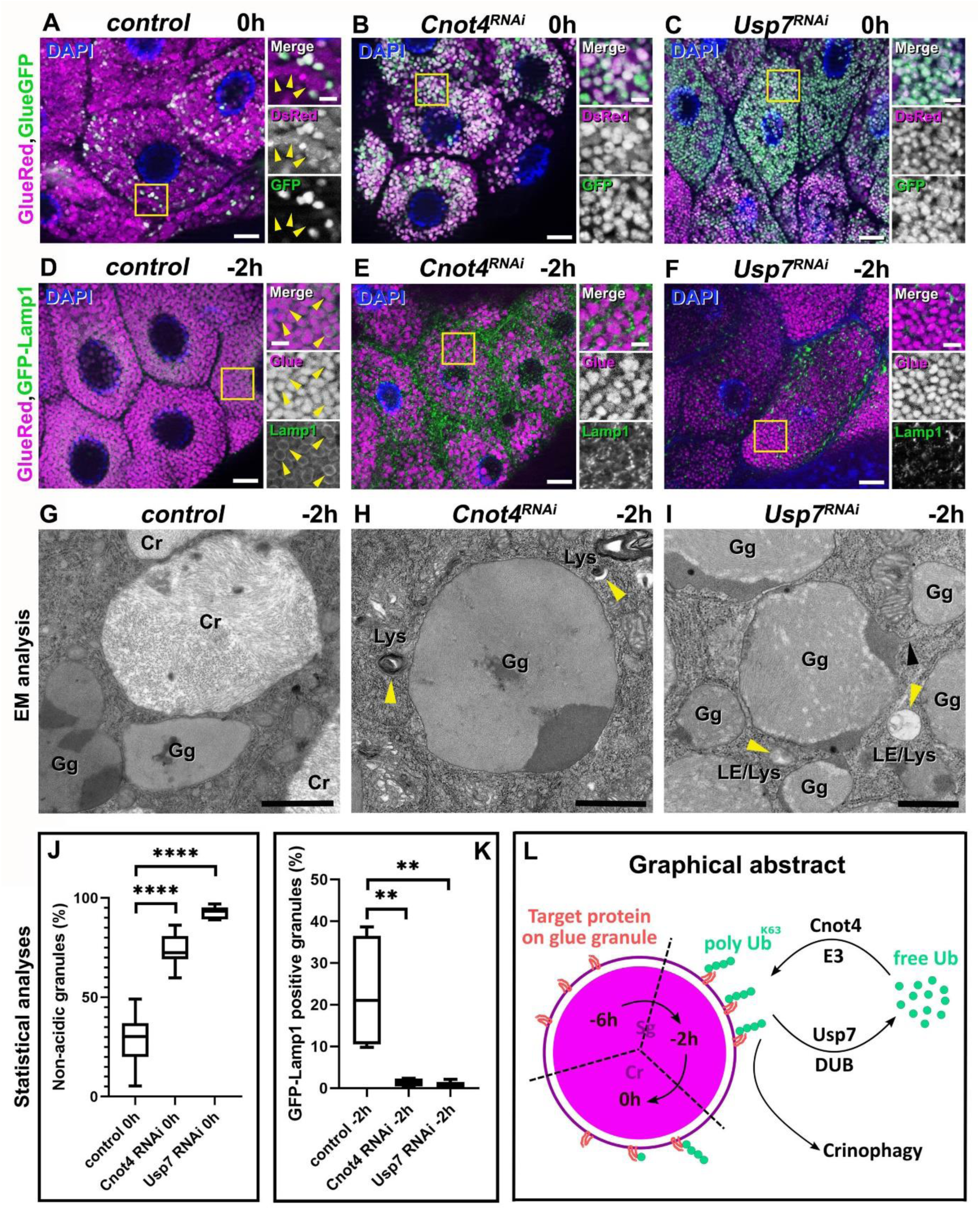
Loss of *Cnot4* or *Usp7* impairs glue granule-lysosome fusions. (**A-C**) GFP is not quenched in glue-containing secretory granules of prepupal (0h RPF) salivary gland cells co-expressing glue-GFP/glue-DsRed reporters and RNA interference for genes in-volved in ubiquitination and deubiquitination of glue granule membranes. Salivary gland-specific knockdown of *Cnot4* (**B**) and *Usp7* (**C**) disrupts the developmentally programmed quenching of GFP fluorescence within glue granules compared to control cells (**A**, yellow arrowheads). (**D-F**) Impaired fusion of glue granules with late endosomes and lysosomes is observed in *Cnot4* and *Usp7* RNAi cells expressing glue-DsRed and GFP-Lamp1 reporters. The formation of GFP-Lamp1 rings around glue-DsRed positive secretory granules (yellow arrowheads) seen in control cells (**D**) is impaired in salivary gland cells undergoing *Cnot4* (**E**) and *Usp7* (**F**) RNAi, indicating a fusion defect between glue granules and endo-lysosomal vesicles. (**G-I**) Ultrastructural analysis confirms secretory granule-late endosome/lysosome fusion defects in salivary gland cells from the late larval stage, upon silencing of *Cnot4* and *Usp7*: intact glue granules persist in *Cnot4* (**H**) and *Usp7* (**I**) knockdown cells. Both intact glue granules and crinosomes containing loose filamentous contents are evident in wild-type gland cells at -2h RPF (**G**). The yellow arrowheads on panels (**H**) and (**I**) indicate the accumulation of late endosomes (LE) and lysosomes (Lys) around intact glue granules (Gg). Additionally, the black arrowhead on panel (**I**) shows the protrusion of a glue granule, which may arise due to impaired recycling of glue granule membrane proteins and lipids in the *Usp7* silenced cells. Quantification of data from panels A-C (**J**) and E-F (**K**) was performed on samples from 6-8 animals using Mann-Whitney tests. P values are **** p<0.0001 (**J**) and ** p<0.0022 (**K**). The model depicts that glue granule designation for crinophagic degradation involves ubiquitin and related enzymes Cnot4 and Usp7 (**L**). Abbreviations: E3: ubiquitin ligase, DUB: Deubiquitinating enzyme, Ub: Ubiquitin, K63 Ub: K63-linked polyubiquitin, Sg: Secretory granule, Cr: Crinosome. The boxed regions in panels (**A–F**) are shown enlarged on the right side of each panel. Magenta and green channels of merged images are shown separately as indicated. Bars: (**A-F**) 20 μm, (**G-H**) 1 μm, (**I**) 500 nm, insets 5 μm.

The degradation of glue granules in the late larval/prepupal salivary gland cells occurs through the direct fusion of residual secretory granules and late endosomes-lysosomes. To investigate the functions of *Cnot4* and *Usp7* in this process, we monitored the appearance of the lysosomal membrane marker GFP-Lamp1 (Lysosome Associated Membrane Protein-1) in the membrane of the DsRed-containing granule granules [11,15,16,42]. Control salivary gland cells from the late larval (-2 h RPF) developmental stage were full of glue granules surrounded by GFP-Lamp1 (**Figure 3D and K**). In contrast, small GFP-Lamp1 positive structures accumulated around the DsRed-containing glue granules instead of forming rings in *Cnot4* and *Usp7* RNAi cells, indicating impaired fusion with late endosomes and lysosomes (**Figure 3E, F and K**). Furthermore, ultrastructural analysis of *Cnot4* and *Usp7* silenced salivary gland cells identified unfused late endosomes and lysosomes near the glue granules, and glue granule content retained an immature morphology compared to the control (**Figure 3G-I**).

Our experiments on glue granule acidification and granule-to-lysosome fusion highlight the important roles of Cnot4 and Usp7 in crinophagy in *Drosophila* salivary gland cells during the late larval and prepupal development. Notably, despite their opposing enzymatic functions of Cnot4 as a ubiquitin ligase and Usp7 as a deubiquitinating enzyme, silencing either of these genes results in similar defects in granule acidification and fusion with lysosomes (**Figure 3A-F and J**, **K**), even though *Cnot4* and *Usp7* RNAi had opposite effects on the ubiquitin positivity of glue granules (**Figure 1B**, **C**, **E**, **F and I**). The striking similarity of crinophagy defects caused by *Cnot4* and *Usp7* knockdown in the salivary gland cells suggests that Cnot4 and Usp7 may function cooperatively during crinophagy and may be part of the same pathway that regulates the degradation of glue granules.

### 9. Ubiquitin regulates the crinophagic degradation of secretory granules

During the intracellular transport of cellular organelles and molecules, these components are precisely delivered to their required locations at the right time. Consequently, these processes necessitate intricate regulatory systems, among which ubiquitin and its regulatory enzymes: ubiquitin ligases and proteases represent key elements (**Figure 3L**). Several vesicular transport pathways rely on ubiquitination including endocytosis, cargo sorting, intraluminal vesicle formation during multivesicular body (MVB) biogenesis, modulation of cytoskeletal components, and autophagic mechanisms [29].

Crinophagy, a non-typical autophagic process, involves the fusion of secretory granules with vesicles of the endo-lysosomal system. The well-established function of this process is the degradation and recycling of obsolete secretory contents [11,12], and recent investigations implicated additional roles for crinophagy and crinophagy-like mechanisms, including antigen generation in pancreatic β cells [43,44], secretory granule quality control in exocrine and endocrine cells [3,7], and possible contribution to the secretory granule maturation [4,6].

This study focused on the degradative role of crinophagy, which targets the unreleased glue granules in the late larval salivary gland cells after the burst of secretion. We found that the unreleased glue granules become ubiquitin-positive, and this molecular pattern contains K63-linked polyubiquitin, the most common ubiquitination signal in vesicular trafficking pathways [26,29]. Furthermore, we also identified the E3 enzyme Cnot4 and the deubiquitinating enzyme Usp7, which may collaborate to facilitate the demolition of glue granules during development. Our immunostaining experiments revealed that Cnot4 and Usp7 localize to the surface of glue granules (**Figure 2D and E**), and *Cnot4*-silenced salivary gland cells showed impaired Usp7 localization to glue granules (**Figure 2F and Figure S2E**) compared to control cells (**Figure 2E and Figure S2E**). These suggest that Cnot4 and Usp7 may operate together (or sequentially) in the regulation of crinophagy.

Our findings support that ubiquitin has a broad role in various secretory granule-lysosome fusion events, including both developmentally programmed and prematurely triggered (in *Rab6*-silenced salivary gland cells) forms of crinophagy. Our research provides new molecular insights into crinophagy, advancing our comprehension of this process under both normal and pathological circumstances.

## MATERIALS AND METHODS

### Fly stocks and work

The following fly stocks were obtained from the Bloomington *Drosophila* Stock Center: *Sgs3 (Glue)-GFP (5884)* [41], *UAS-LifeAct-Ruby (35545)* and *UAS-Rab6^JF02640^ (27490)*. Fly stocks RNAi lines obtained from the Vienna *Drosophila* Resource Centre were: *UAS-Cnot4^GD4410^ (v10850)*, *UAS-Usp7^GD7628^ (v18231)*. Additional fly lines included *UAS-GFP-Ub* (provided by P. Deák, Department of Genetics, University of Szeged, Szeged, Hungary) [31], *UAS-GFP-Lamp1* and *UAS-Vps16A^RNAi^* (provided by H. Krämer, Center for Basic Neuroscience, UT Southwestern Medical Center) [42], *Sgs3 (Glue)-DsRed* (Glue-Red, provided by A. Andres, University of Nevada, Las Vegas, NV) [32], and *fkh-Gal4* [3,11], *UAS-Cnot4* (kindly provided by Mika Rämet, Tampere University, Faculty of Medicine and Health Technology, Finland) [45]. *UAS-Usp7* (kindly provided by Peter Verrijzer and Jan van der Knaap, Erasmus University Medical Center, Rotterdam, The Netherlands) [46].

### Fluorescent microscopy and immunocytochemistry

Salivary glands were dissected from control, mutant, and RNAi animals at the indicated developmental stages, fixed for 5 min in 4% paraformaldehyde in PBS, and covered with PBS/glycerin (9:1) containing DAPI. Ubiquitin (Ub), K63-linked polyubiquitin (UbK63), Cnot4 and Usp7 were detected essentially as described previously (Takáts et al., 2013). In brief, salivary glands were dissected in ice-cold PBS then fixed with 4% formaldehyde in PBTX (0.1% Triton X-100 in PBS for overnight at 4°C). Samples were extensively washed with PBTX (3 × 15 min at RT) and then incubated in blocking solution (5% FCS in PBTX for 30 min at RT). Samples were then incubated with 1. monoclonal mouse anti-Ub (clone A-5: sc366553; Enzo) diluted 1:100, 2. monoclonal rabbit αUbK63 (clone JM09-67; Invitrogen) diluted 1:100, 3. polyclonal rabbit αCnot4 (PA5-101501; Invitrogen) diluted 1:100 and 4. polyclonal rabbit αUsp7 (PA5-34911; Invitrogen) diluted 1:100 in the blocking solution overnight at 4°C. Salivary glands were then washed (3 × 15 min in PBTX at RT) and incubated in blocking solution again for 30 min at RT, followed by incubation with DyLight 488–conjugated goat α–rabbit (SA5-10018; Thermo Fisher Scientific) diluted 1:600 in blocking solution for 3 h at RT. Washing steps were repeated, and samples were mounted with PBS/glycerol (9:1) containing DAPI. Images were taken at RT using a Carl Zeiss AxioImager M2 epifluorescent microscope equipped with an Apotome grid confocal unit and a led lamp, using AxioCam MRm camera Plan-Apochromat 63x NA = 1.4, EC Plan-Neofluar 40x NA = 0.75 objective and processed in Zeiss AxioVision SE64 Rel. 4.9.1 and Adobe Photoshop CS3 Extended.

### Transmission electron microscopy (TEM) and immunogold labeling

Progressive lowering temperature embedding and subsequent immunolabeling were performed as previously described (Lőrincz et al., 2014). In brief, salivary glands from *UAS-Usp7 RNAi* expressing animals were dissected in PBS and fixed with 4% formaldehyde, 0.05% glutaraldehyde, and 0.2% tannic acid in phosphate buffer (PBS; 0.1 M, pH 7.4) overnight at 4°C. Samples were then washed extensively with PBS, and free aldehyde groups were quenched with 50 mM glycine and 50 mM NH_4_Cl in PBS. Salivary glands were then postfixed in 1% uranyl acetate in 0.05 M maleate buffer (3 h at RT). Samples were then dehydrated in a graded series of ethanol as follows: 25% EtOH (10 min, 0°C), 50% EtOH (10 min, 0°C), 70% EtOH (10 min, -20°C), 96% EtOH (20 min, -20°C), and absolute EtOH (2 × 60 min, -20°C). Next, salivary glands were infiltrated with pure LR White (Sigma-Aldrich) containing 2% benzoyl peroxide as catalyst (24 h, -20°C). Curing was performed using a homemade UV chamber (equipped with two 2 × 6-W UV lamps) for 48 h at -20°C. Ultrathin sections (80-90 nm) were cut and collected on nickel grids. All immunoreactions were performed at RT. The following incubations were performed: (1) 5% H_2_O_2_ for 5 min; (2) bi-distilled water for 3 × 5 min; (3) 0.3% NaBH_4_, 50 mM glycine and 50 mM NH_4_Cl in PBS, pH 7.4, for 5 min; (4) PBS for 3 × 5 min; (5) 1% BSA in PBS for 3 × 5 min; (6) 5% FCS and 1% BSA in PBS for 30 min; (7) primary antibody (monoclonal rabbit αUb^K63^ (Invitrogen, clone JM09-67)) diluted 1:25 in 1% BSA in PBS overnight at 4°C; (8) 1% BSA in PBS for 3 × 5 min; (9) 1% BSA in TBS for 3 × 5 min; (10) secondary antibody (10 nm gold-conjugated goat anti-rabbit (Sigma, G-3779)) diluted 1:100 in 1% BSA in TBS for 4 h; (11) TBS for 3 × 5 min; (12) extensive wash with bidistilled water. Sections were viewed in a transmission electron microscope (JEM-1011; JEOL) equipped with a digital camera (Morada; Olympus) using iTEM software 5.1 (Olympus).

### Immunoblotting experiments

Fifty pairs of salivary glands were collected in PBS. Following PBS removal by centrifugation, the glands were resuspended in homogenization buffer (0.2 M sucrose, 10 mM Tris, pH 7.2), snap-frozen in liquid nitrogen, and stored at -80°C. To maintain glue granule integrity during homogenization frozen glands were thawed in a room temperature water bath and immediately transferred to ice (adapted from ref. [47]). Fresh protease inhibitor cocktail was added to the homogenization buffer. The lysate was pre-cleared by incubating 1 ml of homogenate with 25 µl of pre-blocked magnetic agarose beads (Chromotek, bab-20) for 1 hour at 4°C with rotation. Beads were pre-equilibrated with 3 x 500 µl of ice-cold dilution buffer (10 mM Tris/HCl pH 7.5, 150 mM NaCl, 0.5 mM EDTA). The supernatant was collected using a magnetic separator (DynaMag -2) and then incubated with 25 µl of equilibrated GFP-TRAP beads (Chromotek, gtma-20) for 1 hour at 4°C with rotation. After binding, beads were washed 5 x 500 μl of dilution buffer. Bound proteins were eluted by boiling the beads in 2x Laemmli buffer at 95°C for 10 minutes. Input and immunoprecipitated (IP) samples were separated by 10% SDS-PAGE and analyzed by western blot, as previously described [48]. Blots were probed with chicken α-GFP (Aves Labs, GFP-1020, 1:5000), mouse anti-mono- and polyubiquitin (Enzo, BML-PW-88-10-0500, 1:1000), and rabbit anti-DsRed (Sigma, AB356483, 1:3000) primary antibodies. Protein bands were detected using LI-COR fluorescent secondary antibodies (926-32218, 926-33210, and 926-32211, 1:15,000) and visualized on a LI-COR Odyssey CLx scanner using Image Studio software.

### Statistical analysis

Fluorescence structures from original, unmodified single focal planes were quantified manually. Three to six cells were randomly selected for counting from pictures of control and RNAi salivary glands from six to eight animals. In GFP-ubiquitin, GFP-Lamp1 and αUsp7 experiments, GFP-ubiquitin, GFP-Lamp1 and αUsp7 rings around the glue-DsRed granules were counted as double positive. In glue-GFP and glue-DsRed experiments, all of granules were counted and the number of glue-GFP granules were proportioned to the number of all granules. We used GraphPad Prism 8 for data analysis. Mann–Whitney tests were used for comparing two samples. In the box plots, bars show the data ranging between the upper and lower quartiles; median is indicated as a horizontal black line within the box. Whiskers represent the smallest and largest observations. Exact p-values can be found in the figure descriptions.

## ABBREVIATIONS

Atg: Autophagy related
BDSC: Bloomington *Drosophila* Stock Center
CCR4-NOT: Carbon Catabolite Repression 4 - Negative On TATA-less
Cnot4: CCR4-NOT transcription complex, subunit 4
CORVET: class C cORe Vacuole/Endosome Tethering
CP: Cytoplasm
Cr: Crinosome
DsRed: Discosoma (a coral genus) red
*dor*: deep orange
DUB: DeUBiquitinase or DeUBiquitinating enzyme
E3 enzyme: ubiquitin ligase
Gg: Glue granule
HOPS: HOmotypic fusion and Protein Sorting
K63: Lysine 63
Lamp1: Lysosome Associated Membrane Protein-1
LE: Late Endosome
*lt*: light
Lys: Lysosome
MVB: multivesicular body
*Not1*: Negative On TATA-less
*Rcd-1*: Required for cell differentiation 1
RING: Really Interesting New Gene
RNAi: RiboNucleic Acid interference
RPF: Relative to Puparium Formation
Sgs3: Salivary gland secretion 3
Sg: Secretory granule
SNARE: Soluble NSF Attachment Protein Receptor
Syx13: Syntaxin 13
TFEB: Transcription Factor EB
TGN: Trans Golgi Network
*UAS*: Up-stream Activating Sequence
Ub: Ubiquitin
Usp7: Ubiquitin specific protease 7
VDRC: Vienna *Drosophila* Resource Center
Vid: Vacuolar import and degradation
Vps16A: Vacuolar protein sorting 16A.

## ACKNOWLEDGMENTS

We thank Sarolta Pálfia, Ivett Répássy and Ágnes Vinczellér for skillful technical assistance. We also thank our retired colleague Lajos László for the useful discussions about ubiquitin, the fellow worker of HUN-REN Biological Research Centre Szeged Enikő Lakatos for help in data evaluation and our master student Erika Farkas and bachelor student Bori Balatoni for help in basic fly work in the laboratory.

## Author Contributions

T. Csizmadia designed research with input from Péter Lőw and Gábor Juhász. T. Csizmadia, Anna Dósa, Asha Kiran, András Jipa, Hajnalka Laczkó-Dobos and Péter Lőw performed experiments. T. Csizmadia evaluated data. T. Csizmadia and G. Juhász acquired funding, T. Csizmadia, Péter Lőw and Gábor Juhász wrote the paper with comments from all authors.

