## Supplemental material for "Ubiquitination of secretory granules promotes crinophagic degradation in *Drosophila*"

Title

SUPPLEMENTARY MATERIALS

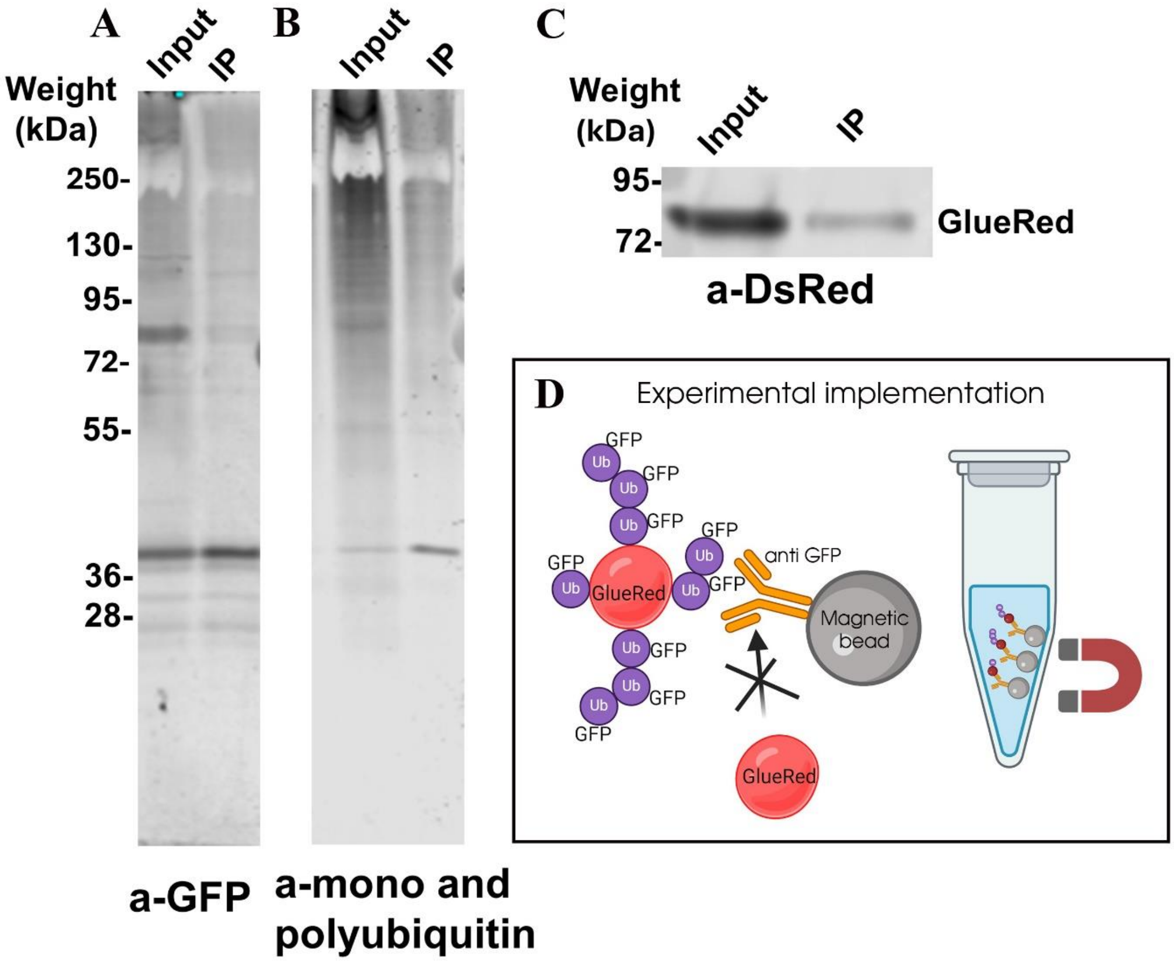


**Figure S1. GFP-trap immunoprecipitation of glue granules from salivary gland cells coexpressing glue-DsRed and GFP tagged ubiquitin (GFP-Ub).** Western blot analysis using GFP (**A**), ubiquitin (recognizing both mono- and polyubiquitin) (**B**) and DsRed (**C**) antibodies confirmed the presence of both proteins (glue-DsRed and ubiquitin) in the input and the immunoprecipitated (IP) fraction. Panel (**D**) shows experimental implementation (generated in BioRender).

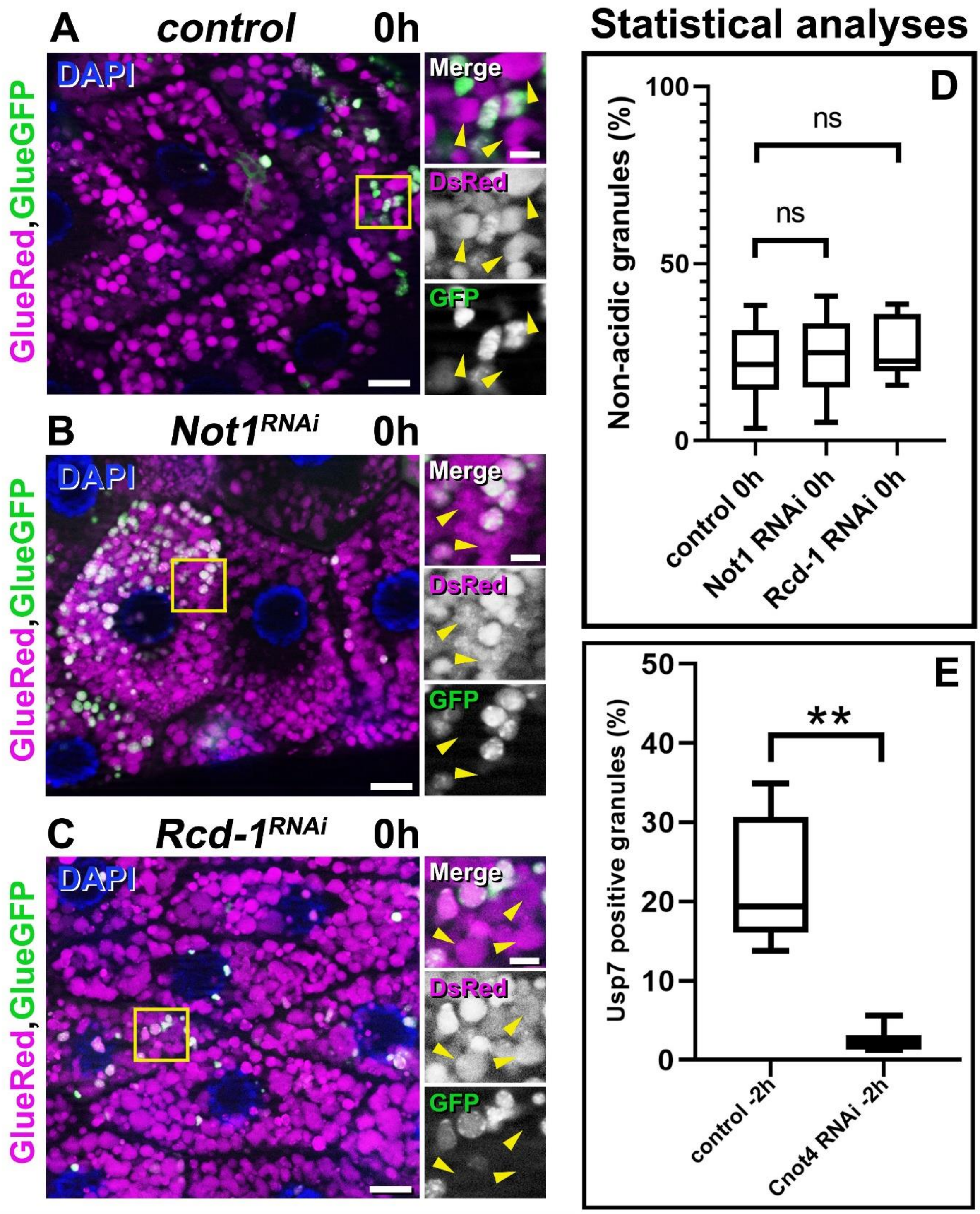


**Figure S2. Silencing genes encoding other subunits of the CCR4-CNOT complex does not influence glue granule acidification**

Prepupal salivary gland cells co-expressing glue-GFP and glue-DsRed reporters and RNA interference for genes encoding subunits of the CCR4-CNOT complex. Salivary gland-specific knockdown of *Not1* (**B**) and *Rcd-1* (**C**) did not disrupt the developmentally programmed quenching of GFP fluorescence within glue granules: these are similar to control cells (**A,** yellow arrowheads). (**D)** Quantification of data from **A-C**, n=6-9 animals, Mann-Whitney tests. P values are (ns) p=0.6842 and (ns) p=0.3527, respectively.

(**E**) Statistical analysis of Usp7 localization on glue granules in late larval (-2 h RPF) salivary gland cells. Quantification of data from **Figure 2E and F**, n=6-8 animals, Mann-Whitney tests, ** $p<0.0022$.

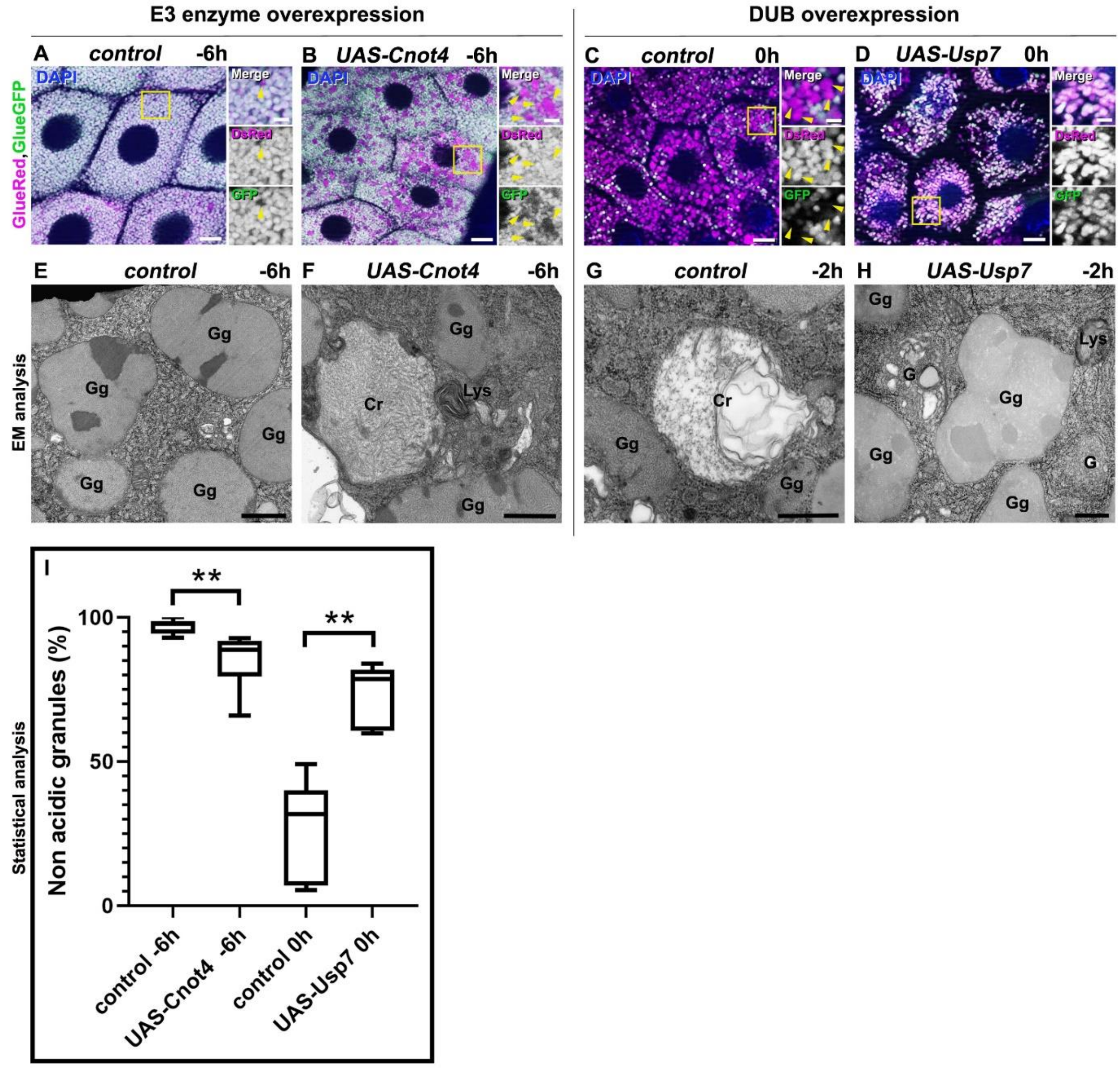


**Figure S3. Overexpression of Cnot4 prematurely induces while Usp7 inhibits developmental crinophagy in salivary gland cells**

(**A-B**) Upon overexpression of Cnot4, acidification of glue granules occurs well before the developmental program in salivary gland cells from wandering animals co-expressing glue-GFP and glue-DsRed reporters. Salivary gland specific overexpression of Cnot4 causes early and intense quenching of GFP fluorescence within crinosomes (**B**, yellow arrowheads), compared to the control cells (**A**). (**C-D**) Usp7 overexpression impairs the acidification of glue-containing secretory granules in prepupal (0 h RPF) salivary gland cells co-expressing glue-GFP and glue-DsRed reporters. Salivary gland-specific overexpression of Usp7 (**D**) disrupts the developmentally programmed quenching of GFP fluorescence within glue granules, compared to control cells (**C**, yellow arrowheads. (**E-H**) Ultrastructural analysis of salivary gland cells overexpressing Cnot4 (**F**) and Usp7 (**H**). Intense crinosome formation is observed in Cnot4 overexpressing salivary gland cells from wandering animals (**F**), unlike in the control (**E**). Intact glue granules persist in Usp7 (**H**) overexpressing cells. Both intact glue granules and crinosomes containing loose filamentous contents are evident in wild-type gland cells at -2h RPF (**G**). Quantification of data from panels A-D (**I**) was performed on samples from 6-8 animals using Mann-Whitney tests. P values are ** $p<0.0022$ (**I**). Abbreviations: Gg: Glue granule, Cr: Crinosome, Lys: lysosome, G: Golgi. The boxed regions in panels (**A–D**) are shown enlarged on the right side of each panel. Magenta and green channels of merged images are shown separately as indicated. Bars: (**A-D**) 20 µm, insets 5 µm, (**E**) 2 µm, (**F and H**) 500 nm, (**G**) 1.5 µm.

**Table S1.** Results of the RNAi screen of the *Drosophila* E3 enzyme encoding genes.

| Potential *Drosophila* E3 Ub ligases examined in this study | | | | | | |
|---|---|---|---|---|---|---|
| ***Type of E3 enzyme/domain*** | ***Number*** | ***Name or CG number*** | ***RNAi line*** | ***Source*** | ***Crinophagy status*** | ***Other comment*** |
| **Goliath type** | 1 | gol | 2679R-2 | NigFly | no effect | — |
| | 2 | gzl | 10277R-3 | NigFly | no effect | — |
| **HECT type** | 3 | CG3356 | 3356R-1 | NigFly | no effect | — |
| | 4 | CG4238 | v41982 | VDRC | no effect | — |
| | 5 | CG5087 | 5087R-3 | NigFly | no effect | — |
| | 6 | CG42797 | 3003Ra-4 | NigFly | no effect | — |
| | 7 | ctrip | HMS00322 | NigFly | no effect | — |
| | 8 | HERC2 | 11734R-3 | NigFly | no effect | — |
| | 9 | Herc4 | v37220 | VDRC | no effect | — |
| | 10 | HUWE1 | 8184R-1 | NigFly | no effect | — |
| | 11 | hyd | 9484R-2 | NigFly | no effect | small salivary glands and granules |
| | 12 | Nedd4 | 7555R-3 | NigFly | no effect | — |
| | 13 | Smurf | 4943R-2 | NigFly | no effect | — |
| | 14 | Su(dx) | 4244R-1 | NigFly | no granules | very small salivary gland |
| | 15 | Ube3a | 31972 | BDSC | no effect | — |
| | 16 | Ufd4 | 5604R-3 | NigFly | persisting GFP | — |
| **IAP type** | 17 | Diap1 | 12284R-2 | NigFly | no effect | — |
| | 18 | Diap2 | 8293R-1 | NigFly | no effect | — |
| **Kcmf1 type** | 19 | CG3526 | v26214 | VDRC | no effect | — |
| | 20 | CG15286 | 15286R-3 | NigFly | no effect | — |
| | 21 | CG31642 | HMJ21895 | NigFly | no effect | — |
| | 22 | CG31835 | 31835R-2 | NigFly | no effect | — |
| | 23 | CG42585 | HMJ23964 | NigFly | no effect | — |
| | 24 | Kcmf1 | 11984R-2 | NigFly | no granules | — |
| **Other type** | 25 | CG3894 | 42618 | BDSC | no effect | — |
| | 26 | CG7326 | 7326R-1 | NigFly | no effect | — |
| | 27 | CG13994 | 13994R-2 | NigFly | no effect | — |
| | 28 | CG14646 | 14646R-4 | NigFly | persisting GFP | variable effect |
| | 29 | I-3 | v45386 | VDRC | no effect | — |
| | 30 | Neurl4 | 44094 | BDSC | no effect | — |
| | 31 | poe | 14472R-1 | NigFly | no effect | — |
| **Other Ring types** | 32 | Bre1 | HMJ22277 | NigFly | no effect | — |
| | 33 | Cbl | 7037R-1 | NigFly | no effect | — |
| | 34 | CG1317 | HMJ21301 | NigFly | no effect | — |
| | 35 | CG1909 | HMC02933 | NigFly | no effect | — |
| | 36 | CG2617 | 2617R-1 | NigFly | no effect | — |
| | 37 | CG2617 | HMJ21915 | NigFly | no effect | — |
| | 38 | CG2681 | 2681R-1 | NigFly | no effect | — |

| | | | | | | |
|---|---|---|---|---|---|---|
| | 39 | CG2926 | 2926R-2 | NigFly | persisting GFP | very small gland |
| | 40 | CG2991 | 2991R-3 | NigFly | no effect | — |
| | 41 | CG4080 | 4080R-2 | NigFly | no effect | — |
| | 42 | CG4325 | v34829 | VDRC | no effect | — |
| | 43 | CG4813 | 4813R-1 | NigFly | persisting GFP | small granules |
| | 44 | CG5071 | 5071R-1 | NigFly | no effect | — |
| | 45 | CG5334 | 5334R-1 | NigFly | no effect | — |
| | 46 | CG5347 | 5347R-1 | NigFly | persisting GFP | small granules |
| | 47 | CG5382 | v101394 | VDRC | no effect | — |
| | 48 | CG5555 | 5555R-2 | NigFly | persisting GFP | — |
| | 49 | CG6752 | 6752R-1 | NigFly | no effect | — |
| | 50 | CG6923 | 6923Ra-3 | NigFly | no effect | — |
| | 51 | CG7376 | v35222 | VDRC | no effect | — |
| | 52 | CG7694 | 7694R-1 | NigFly | no effect | — |
| | 53 | CG8141 | HMJ23674 | NigFly | no effect | — |
| | 54 | CG8910 | 8910R-3 | NigFly | persisting GFP | variable effect, small granules |
| | 55 | CG8974 | not available | — | — | — |
| | 56 | CG9014 | 9014R-1 | NigFly | no effect | — |
| | 57 | CG9855 | HMJ23874 | NigFly | no effect | — |
| | 58 | CG9941 | v29596 | VDRC | no effect | — |
| | 59 | CG10761 | v5474 | VDRC | no effect | — |
| | 60 | CG10916 | HMJ23868 | NigFly | no effect | — |
| | 61 | CG11360 | 11360R-4 | NigFly | no effect | — |
| | 62 | CG11414 | 11414R-3 | NigFly | no effect | — |
| | 63 | CG12099 | v18734 | VDRC | no effect | — |
| | 64 | CG12477 | v31944 | VDRC | no effect | — |
| | 65 | CG13025 | 13025R-2 | NigFly | no effect | — |
| | 66 | CG13344 | 13344R-2 | NigFly | no effect | — |
| | 67 | CG13442 | HMS01518 | NigFly | no effect | — |
| | 68 | CG13481 | v103527 | VDRC | no effect | — |
| | 69 | CG13605 | v105112 | VDRC | no effect | — |
| | 70 | CG14435 | 14435R-4 | NigFly | no granules | very small salivary gland |
| | 71 | CG14983 | HMJ23801 | NigFly | no effect | — |
| | 72 | CG15011 | 15011R-3 | NigFly | no effect | — |
| | 73 | CG15141 | 15141R-1 | NigFly | no effect | — |
| | 74 | CG15814 | v30430 | VDRC | no effect | — |
| | 75 | CG16781 | v7020 | VDRC | no effect | — |
| | 76 | CG17019 | 17019R-6 | NigFly | no effect | — |
| | 77 | CG17048 | v8780 | VDRC | no effect | — |
| | 78 | CG17260 | 17260R-1 | NigFly | no effect | — |
| | 79 | CG17329 | v19171 | VDRC | no effect | — |
| | 80 | CG17717 | 17717R-2 | NigFly | no effect | — |
| | 81 | CG17721 | v6036 | VDRC | no effect | — |

| | | | | | | |
|---|---|---|---|---|---|---|
| | 82 | CG17991 | 17991R-1 | NigFly | no effect | — |
| | 83 | CG31807 | 31807R-2 | NigFly | no effect | — |
| | 84 | CG32369 | 64028 | BDSC | no effect | — |
| | 85 | CG32581 | not available | — | — | — |
| | 86 | CG32847 | v48423 | VDRC | no effect | — |
| | 87 | CG32850 | HMJ22085 | NigFly | no effect | — |
| | 88 | CG33552 | HMJ23596 | NigFly | no effect | — |
| | 89 | CG34289 | HMJ21306 | NigFly | no effect | — |
| | 90 | CG34308 | not available | — | — | — |
| | 91 | CG34375 | 13835R-2 | NigFly | no effect | — |
| | 92 | Cnot4 | 31716R-1 | NigFly | persisting GFP | — |
| | 93 | Cnot4 | v10850 | VDRC | persisting GFP | — |
| | 94 | d4 | 2682R-1 | NigFly | persisting GFP | — |
| | 95 | dgrn | 10981R-3 | NigFly | no effect | — |
| | 96 | dnr1 | 12489R-1 | NigFly | persisting GFP | small granules |
| | 97 | dor | 3093R-2 | NigFly | persisting GFP | — |
| | 98 | dx | 3929R-1 | NigFly | no effect | — |
| | 99 | elfless | 15150R-1 | NigFly | no effect | — |
| | 100 | elgi | 17033R-3 | NigFly | no effect | — |
| | 101 | Fancl | 12812R-1 | NigFly | no effect | — |
| | 102 | Ltn1 | 9268R-3 | NigFly | no effect | — |
| | 103 | Mat1 | 7614R-1 | NigFly | no effect | — |
| | 104 | mdlc | 4973R-1 | NigFly | no effect | — |
| | 105 | Hakai | 2LG-0654 | NigFly | no effect | — |
| | 106 | hiw | 28031 | BDSC | no effect | — |
| | 107 | Iru | 11982R-3 | NigFly | no effect | — |
| | 108 | CG4195 | 4195R-1 | NigFly | no effect | — |
| | 109 | ImgA | 2LG-0313 | NigFly | no effect | — |
| | 110 | Lpt | 25994 | BDSC | no effect | — |
| | 111 | lt | 18028R-2 | NigFly | persisting GFP | — |
| | 112 | Mi-2 | HMS00301 | NigFly | no effect | — |
| | 113 | mib1 | 27320 | BDSC | no effect | — |
| | 114 | mib2 | HMJ21843 | NigFly | no effect | — |
| | 115 | Mkrn1 | v34373 | VDRC | no effect | — |
| | 116 | msl-2 | 31627 | BDSC | no effect | — |
| | 117 | Mul1 | 1134R-2 | NigFly | no effect | — |
| | 118 | Mura | GL00121 | NigFly | no effect | — |
| | 119 | neur | 11988R-1 | NigFly | no effect | — |
| | 120 | nopo | v22013 | VDRC | no effect | — |
| | 121 | Nse1 | 2LG-0755 | NigFly | no effect | — |
| | 122 | Pex2 | 7081R-2 | NigFly | no effect | — |
| | 123 | Pex10 | v46613 | VDRC | no effect | — |
| | 124 | Pex12 | 3639R-1 | NigFly | no effect | — |
| | 125 | Pli | 5212R-3 | NigFly | no effect | — |
| | 126 | POSH | 4909R-1 | NigFly | no effect | — |

| | | | | | | |
|---|---|---|---|---|---|---|
| | 127 | Psc | 3886R-1 | NigFly | no effect | — |
| | 128 | qin | HMJ21020 | NigFly | no effect | — |
| | 129 | Rbpn-5 | 4030R-3 | NigFly | no effect | — |
| | 130 | Rchy1 | HMJ22014 | NigFly | no effect | — |
| | 131 | Rnf146 | 8786R-4 | NigFly | no effect | — |
| | 132 | roq | HMJ21890 | NigFly | no effect | — |
| | 133 | Sce | 5595R-1 | NigFly | persisting GFP | very few and small granules |
| | 134 | sina | 9949R-1 | NigFly | no effect | — |
| | 135 | sinah | 13030R-2 | NigFly | no effect | — |
| | 136 | sip3 | 1937R-3 | NigFly | no effect | — |
| | 137 | snama | 3231R-3 | NigFly | no effect | — |
| | 138 | snky | HMJ21388 | NigFly | no effect | — |
| | 139 | stc | HMS02768 | NigFly | no effect | — |
| | 140 | Su(z)2 | HMS00281 | NigFly | no effect | — |
| | 141 | Topors | 15104R-2 | NigFly | no effect | — |
| | 142 | Traf6 | 10961R-1 | NigFly | no effect | — |
| | 143 | Trc8 | v4449 | VDRC | no effect | — |
| | 144 | trx | HMS00580 | NigFly | no effect | — |
| | 145 | Ubr1 | 9086R-3 | NigFly | no effect | — |
| | 146 | Ubr3 | v22901 | VDRC | no effect | — |
| | 147 | Unk | 57026 | BDSC | no effect | — |
| | 148 | Vps8 | 10144R-2 | NigFly | no effect | — |
| | 149 | Vps11 | v24731 | VDRC | persisting GFP | — |
| | 150 | ari-1 | 29416 | BDSC | no effect | — |
| **Ring btw Ring** | 151 | ari-2 | 5709R-1 | NigFly | no effect | — |
| | 152 | CG12362 | 12362R-1 | NigFly | no effect | — |
| | 153 | CG33144 | 64033 | BDSC | no effect | — |
| | 154 | LUBEL | 11321R-3 | NigFly | no effect | — |
| | 155 | park | 10523R-2 | NigFly | no effect | — |
| **Roc types** | 156 | Roc1a | HMS00353 | NigFly | no effect | — |
| | 157 | Roc1b | 31067 | BDSC | no effect | — |
| | 158 | Roc2 | v28103 | VDRC | no effect | — |
| **Trim types** | 159 | bon | 27047 | BDSC | no effect | — |
| | 160 | CG8419 | v107626 | VDRC | no effect | — |
| | 161 | mei-P26 | 12218R-1 | NigFly | no effect | — |
| | 162 | tn | HMS02508 | NigFly | no effect | — |
| | 163 | Trim9 | 31721R-2 | NigFly | no effect | — |
| **U-box types** | 164 | CG2218 | 2218R-4 | NigFly | no effect | — |
| | 165 | CG6197 | 6179R-1 | NigFly | no effect | — |
| | 166 | CG7747 | v44854 | VDRC | no effect | — |
| | 167 | CG9934 | HMJ21044 | NigFly | no effect | — |
| | 168 | CG11070 | v110416 | VDRC | no effect | — |
| | 169 | Prp19 | 5519R-1 | NigFly | no effect | — |
| | 170 | STUB1 | 5203R-3 | NigFly | no effect | — |

**Table S2.** Results of the RNAi screen of the *Drosophila* DUB encoding genes.

| Examined *Drosophila* deubiquitinating enzymes | | | | | | |
|---|---|---|---|---|---|---|
| ***Type of Deubiquitinating enzyme*** | ***Number*** | ***Name or CG number*** | ***Used stock*** | ***Source*** | ***Crinophagy status*** | ***Other comment*** |
| **UCH DEUBIQUITINASES** | 1 | Uch-L5R | 1950R-1 | NigFly | no effect | — |
| | 2 | Uch-L5 | v103481 | VDRC | no effect | — |
| | 3 | Uch | v26468 | VDRC | no effect | — |
| | 4 | caly | v107757 | VDRC | no effect | — |
| **JAMM DEUBIQUITINASES** | 5 | CG2224 | v110286 | VDRC | no effect | — |
| | 6 | CG4751 | v45530 | VDRC | persisting GFP | variable effect |
| | 7 | Rpn11 | v19272 | VDRC | no granules | very small salivary gland |
| | 8 | eIF3f1 | v101465 | VDRC | no effect | — |
| | 9 | eIF3f2 | v15506 | VDRC | no effect | — |
| | 10 | eIF3h | v106189 | VDRC | no effect | — |
| **USP DEUBIQUITINASES** | 11 | CYLD | v15340 | VDRC | persisting GFP | small granules |
| | 12 | DUBAI | v28960 | VDRC | persisting GFP | — |
| | 13 | faf | v107716 | VDRC | no effect | — |
| | 14 | not | v45775 | VDRC | no effect | — |
| | 15 | puf | v27517 | VDRC | no effect | — |
| | 16 | scny | 27558 | BDSC | no effect | — |
| | 17 | Usp1 | 100992 | VDRC | no effect | — |
| | 18 | Usp2 | v104382 | VDRC | no effect | — |
| | 19 | Usp5 | 12082R-2 | NigFly | persisting GFP | very few and small granules |
| | 20 | Usp7 | v18231 | VDRC | persisting GFP | — |
| | 21 | Usp8 | v107623 | VDRC | no effect | — |
| | 22 | Usp10 | v37859 | VDRC | no effect | — |
| | 23 | Usp12-46 | v27799 | VDRC | no effect | — |
| | 24 | Usp14 | v110227 | VDRC | persisting GFP | variable effect |
| | 25 | Usp15-31 | v103553 | VDRC | no effect | — |
| | 26 | Usp16-45 | v41976 | VDRC | persisting GFP | — |
| | 27 | Usp20-33 | v42609 | VDRC | no effect | — |
| | 28 | Usp30 | v110616 | VDRC | no effect | — |
| | 29 | Usp32 | v18981 | VDRC | no effect | — |
| | 30 | Usp39 | v47664 | VDRC | no effect | — |
| | 31 | Usp47 | v103743 | VDRC | no granules | — |
| **SERINE-TYPE OTU DEUBIQUITINASES** | 36 | CG3251 | v100532 | VDRC | no effect | very few crinosomes |

| | | | | | | |
|---|---|---|---|---|---|---|
| | 37 | otu | v108845 | VDRC | persisting GFP | very few and small granules |
| CYSTEINE-TYPE OTU DEUBIQUITINASES | 38 | CG4968 | 35615 | BDSC | persisting GFP | very few and small granules |
| | 39 | Duba | v109912 | VDRC | no effect | — |
| | 40 | Otud6 | v105469 | VDRC | no effect | — |
| | 41 | Yod1 | v21894 | VDRC | no effect | — |
| | 42 | trbd | v24030 | VDRC | persisting GFP | — |
| OTHERS | 45 | CSN5 | 28732 | BDSC | no effect | — |
| | 46 | CSN6 | v105385 | VDRC | no effect | — |
| | 47 | Josd | v7113 | VDRC | no effect | — |
| | 48 | Rpn8 | v108573 | VDRC | persisting GFP | — |
